# In vitro characterization of human articular chondrocytes and chondroprogenitors derived from normal and osteoarthritic knee joints

**DOI:** 10.1101/440107

**Authors:** Elizabeth Vinod, Upasana Kachroo, Solomon Sathishkumar, P.R.J.V.C Boopalan

## Abstract

**Objective:** Cell based therapy optimization is constantly underway since regeneration of genuine hyaline cartilage is under par. Although single source derivation of chondrocytes and chondroprogenitors is advantageous, lack of a characteristic differentiating marker obscures clear identification of either cell type which is essential to create a biological profile and is also required to assess cell type superiority for cartilage repair. This study was the first attempt where characterization was performed on the two cell populations derived from the same human articular cartilage samples.

**Design:** Cells obtained from normal/osteoarthritic knee joints were expanded in culture (up to passage 10). Characterization studies was performed using flow cytometry, gene expression was studied using RT-PCR, growth kinetics and tri-lineage differentiation was also studied to construct a better biological profile of chondroprogenitors as well as chondrocytes.

**Results and conclusions:** Our results suggest that sorting based on CD34(-), CD166(+) and CD146(+), instead of isolation using fibronectin adhesion assay (based on CD49e+/CD29+), would yield a population of cells primarily composed of chondroprogenitors which when derived from normal as opposed to osteoarthritic cartilage, could provide translatable results in terms of enhanced chondrogenesis and reduced hypertrophy; both indispensable for the field of cartilage regeneration.

## Introduction

Cell-based therapy forms the mainstay of treatment for cartilage afflictions like osteoarthritis (OA) and osteo-chondral defects currently (1). The main contenders in this field which have been used as either stand-alone substitutes, as co-cultures or in combination with scaffolds and growth factors are cartilage derived chondrocytes and mesenchymal stem cells(MSCs) (2,3). Although MSCs (due to inherent multipotency and high replicative potential) (4)and chondrocytes (tissue nativity making them safe for use) (5) are promising candidates; their current usage still warrants optimization. It has been reported that MSCs exhibit a tendency for increased osteogenesis in-vitro (6,7) and fibrocartilage formation *in vivo* (8–10). Similarly, the limitations that affect chondrocyte use (Autologous chondrocyte implantation) are graft hypertrophy and mixed fibro-hyaline formation (11,12). Moreover, chondrocytes require expansion *in- vitro,* since cell yield postharvest is too low to meet demands of direct implantation. This raises another conflict since there does not appear to be a consensus on chondrocyte behavior in culture. There is evidence to show that with increased time in culture, chondrocytes lose their phenotype and show higher expression of markers for hypertrophy thereby reducing their efficiency for optimal cartilage repair(13,14). However, there are also report which demonstrate that chondrocytes exhibit positive stem cell markers in culture(15,16).

Continued search for an optimal cell source led to a potential cell type residing within the superficial layer of cartilage. Isolated by fibronectin adhesion assay, articular cartilage derived chondroprogenitors (CPs) have been classified as MSCs since they demonstrate similar marker profile (Notch-1 signaling proteins, STRO-1, CD90 etc.), high replicative potential, high telomerase activity, and low expression of hypertrophy markers (17–21). Since these cells are native to cartilage and possess progenitor like properties, they appear to be suitable for cartilage repair and inherently primed for chondrogenesis.

Although there are established protocols for isolation of pure populations of CPs (22) and chondrocytes (23) classical differentiating markers between the two cell populations have not been established. Our primary objective was to compare the cell types and evaluate differences in their biological characteristics. Flow cytometric analysis was performed to look for surface marker expression. Reverse Transcription Polymerase Chain Reaction (RT-PCR) was done for assessing markers of chondrogenesis and hypertrophy. The cells were subjected to tri-lineage differentiation to check for multipotency. Cumulative population doubling was done to compare replicative potential of the two cell types. Since CPs have been categorized as MSCs, the first category of surface markers considered for comparison included markers of positive expression: CD105, CD73, CD90, CD106(24,25) and markers of negative expression: CD34, CD45 and CD14(26). The second category for comparison included markers considered to be expressed specifically by chondrocytes: CD54(25) and CD44(27). The final category for comparison included CD markers which are reported to be expressed by cells exhibiting enhanced chondrogenic potential: CD9(28), CD29(29), CD151(30), CD49e(22,28), CD166(31) and CD146(32). To differentiate chondrocytes and CPs on the basis of their chondrogenic potential and tendency for hypertrophy, mRNA expression for markers of chondrogenesis (Collagen type II, Aggrecan and SOX9) and of hypertrophy (Collagen type I, Collagen type X, RUNX2 and MMP-13) was analyzed(33).

Since availability of OA cartilage is comparatively more than normal cartilage, and a dearth of knowledge exists regarding effect of disease on cellular phenotype (34), our second objective was to assess if OA differentially affects the cell populations under consideration, therefore cell samples isolated from normal and OA human cartilage were compared. Our final objective was to study the effects of prolonged time in culture on the cell populations. This would afford additional information about chondrocyte behavior in culture and proposed potency of CPs.

## Materials and Methods

### Study design

The study protocol was approved by the Institutional Review board and carried out in accordance to the guidelines laid down by the Ethics Committee. Human articular cartilage was harvested from three normal (mean age: 22±4yrs) and OA (mean age: 63±7yrs) knee joints. Cartilage was obtained from OA patients undergoing knee replacement surgery and from patients undergoing post-trauma above-knee amputations. Written informed consent as per ethical guidelines was obtained prior to sample collection.

CPs and chondrocytes were isolated from superficial layer/full depth articular cartilage from normal or OA knee joints. Both cell types were then cultured to passage (p) 10. Cells at different time points in culture, namely p0, p1, p2, p3, p4, p5, p7 and p10 were characterized using FACS for the three groups of surface markers mentioned before. Normal and OA cartilage derived CPs and chondrocytes from p0 and p5 were compared for chondrogenic and hypertrophy markers by subjecting them to RT PCR analysis. CPs and chondrocytes from p2 cultures were subjected to tri-lineage differentiation (adipogenic, osteogenic and chondrogenic differentiation). In all, this study contained 4 cell groups for comparison i.e. chondrocytes from normal cartilage, chondrocytes from OA cartilage, CPs from normal cartilage and CPs from OA cartilage (Fig 1). Growth kinetics between the 4 groups was compared at each passage for estimation of cumulative population doubling.

**Fig 1:**
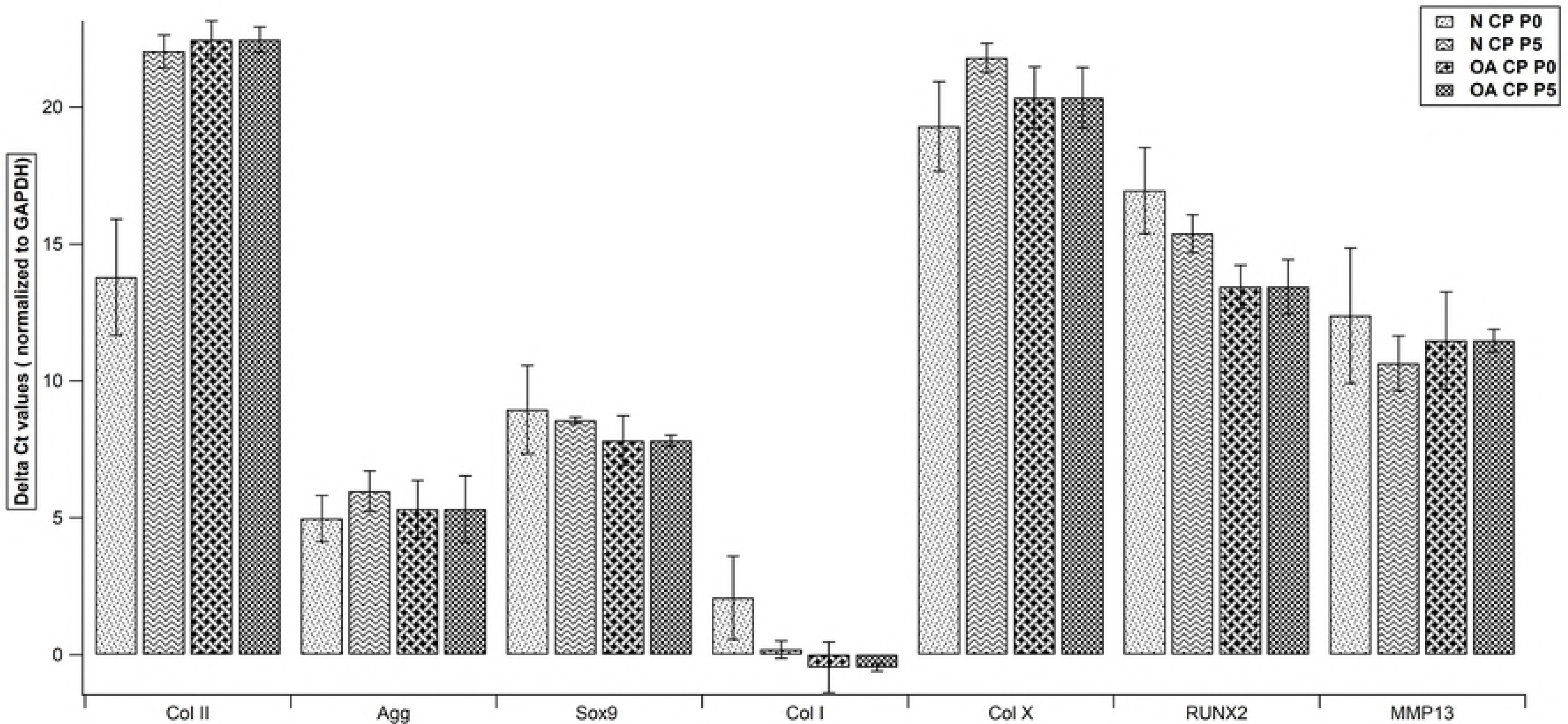
Study algorithm depicting the four groups used for comparison. Samples were taken from three donors in each group. N: normal OA: osteoarthritic, C: chondrocytes, CP: chondroprogenitors, p: passage, CD: cluster of differentiation, SOX9:(sex determining region Y)-box 9, RUNX2: Runt-related transcription factor-2 and MMP-13: metalloproteinase-13.

### Isolation and culture of chondrocytes

Full depth cartilage was harvested from normal and OA knee joints and shavings were washed in phosphate buffered saline (PBS). Overnight cellular digestion was performed using Dulbecco’s modified Eagles medium (DMEM-F12-Himedia) containing 0.15% collagenase type II (Worthington) under standard conditions (23). Following digestion, cells were suspended in medium and cell count was performed. Chondrocytes were loaded at a concentration of 10,000 cells/cm^2^ and expanded to p10 with DMEM/F12 containing 10% fetal calf serum (FCS- Gibco), ascorbic acid 62 μg/ml (Sigma), L-glutamine 2.5mM/L (Sigma), penicillin-streptomycin 100 IU/ml (Gibco) and amphotericin-B 2 μg/ml (Gibco). Medium was changed once every three days (Fig 2). At a confluence of 85-90%, cell harvest was carried out using 0.25% Trypsin-EDTA (Gibco).

**Fig 2:**
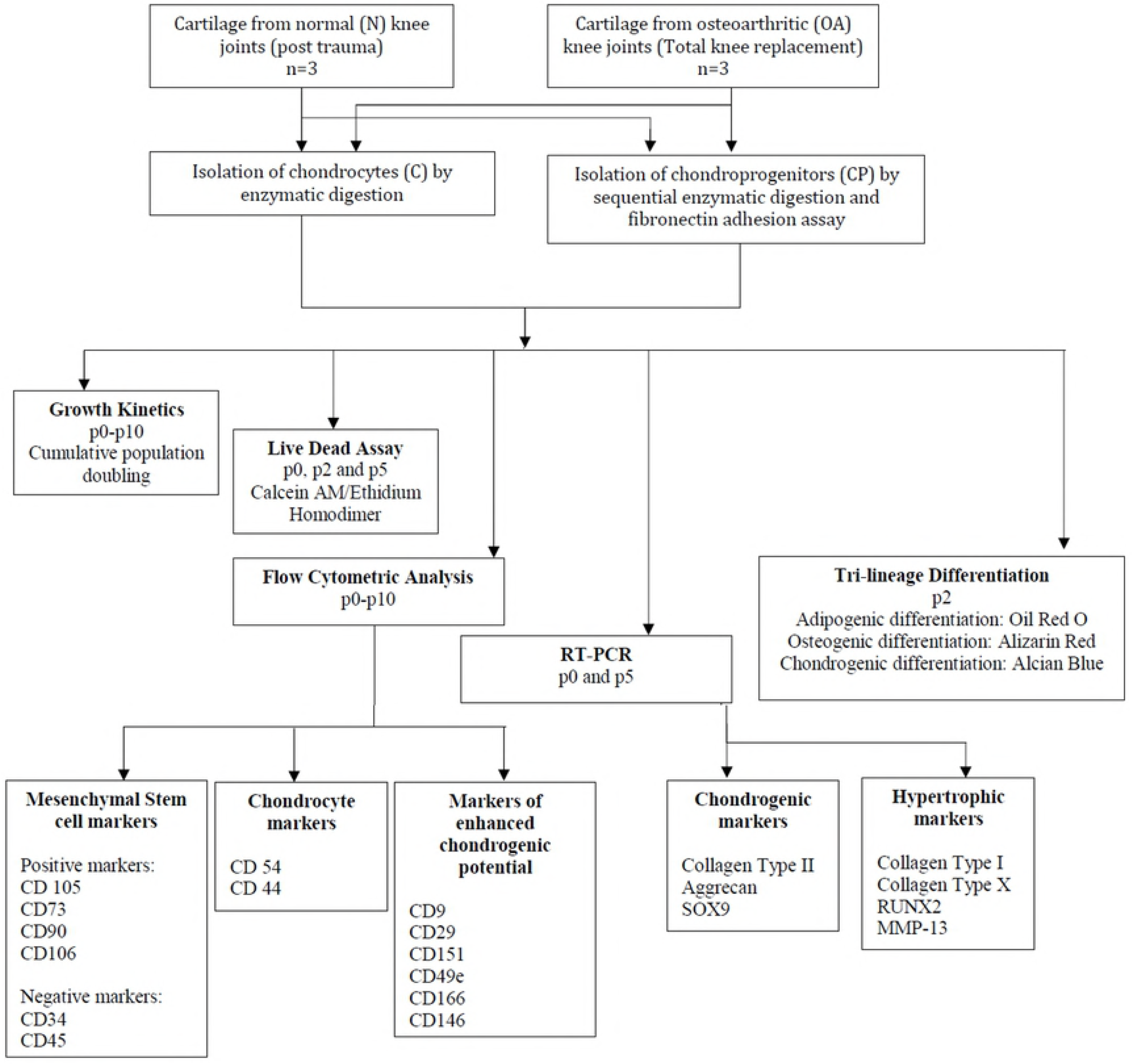
Phase contrast and live dead assay. Upper panel: Phase contrast images of all groups at p0, p2 and p5(10X). Lower panel: Live dead assay images of all groups at sub-confluence showing Calcein AM (live cells-green fluorescence) and Ethidium homodimer (dead cells-red fluorescence) staining at p0, p2 and p5(10X).

### Isolation and culture of articular cartilage derived chondroprogenitors (CPs)

CPs were isolated from the superficial zone of cartilage (same source as used for chondrocytes). Shavings were subjected to sequential overnight enzymatic digestion with 0.2% pronase (Roche) and 0.04% collagenase type II (Worthington) to obtain individual chondrocytes. These were subjected to differential adhesion on fibronectin (Sigma-10μg/ml in PBS containing 1mM CaCl_2_ and 1mM of MgCl_2_) pre-coated plates for 20 min in DMEM containing 10% FCS at a concentration of 700 cells/ml. Post incubation, excess media and non-adherent cells were removed and replaced with standard growth media [DMEM-F12-Glutamax (Himedia) plus ascorbic acid 62 μg/ml(Sigma), L-glutamine 2.5mM/L(Sigma), penicillin-streptomycin 100IU/ml(Gibco) and amphotericin-B 2 μg/ml(Gibco)]. Adherent cells were maintained at standard culture conditions for 10 to 12 days to obtain colonies of >32 cells known as CP clones (Fig 3). Clones were isolated and re-plated at a ratio of 1 clone per 5 cm^2^. Further expansion of enriched polyclonal CPs to p10 was done as per protocol described by Rebecca et al(17) (Fig 2). In brief, medium used was a mixture of Glutamax DMEM-F12 containing 10% FCS supplemented with ascorbic acid 62 μg/ml, L-glutamine 2.5mM/L, penicillin-streptomycin 100 IU/ml, amphotericin-B 2 μg/ml, transforming growth factor beta2 (TGFβ2) 1ng/ml (human-recombinant, Biovision) and fibroblastic growth factor FGF2 5ng/ml (human-recombinant, Biovision). Medium was changed once every three days.

**Fig 3:**
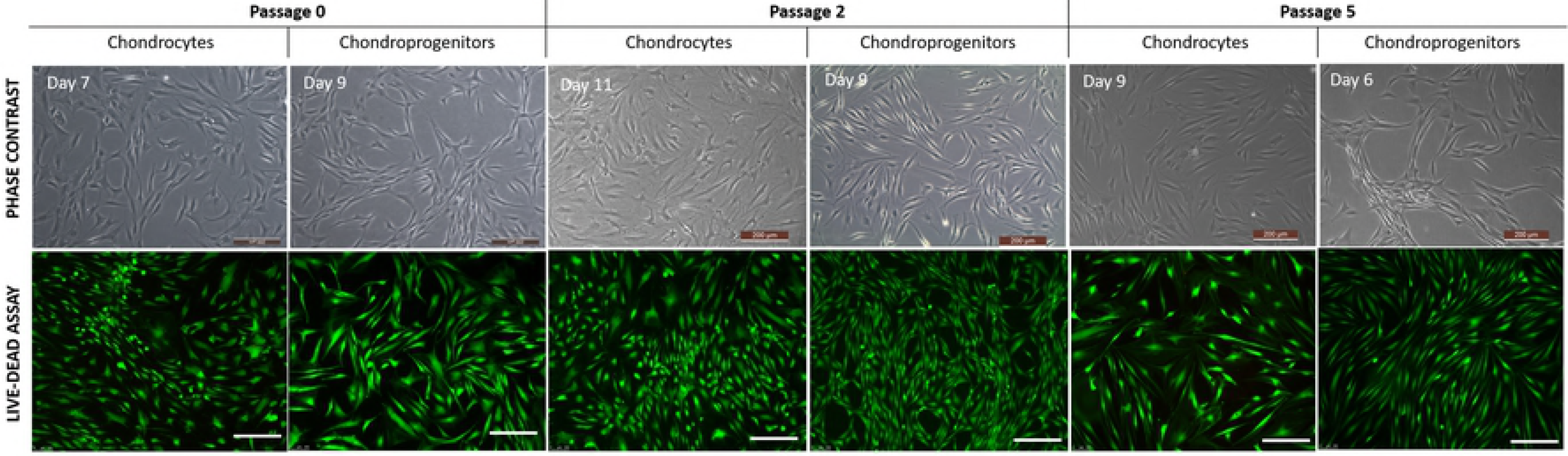
Representative clonally derived human articular chondroprogenitor cells (Fibronectin differential adhesion). A) A clone at day 5 forming a loose cluster of few cells (40X). B) Clone cluster of less than 32 cells at day 8 (10X). C) Clone cluster of more than 40 cells at day 10(10X). D) Live dead assay using Calcein AM/Ethidium homodimer of a clone at day 13(10X).

### Population doubling (PD)

Chondrocytes and CPs were expanded in monolayer cultures and passaged when 85-90% confluent. Cell count was performed using a Neubauer chamber and Population Doubling for all the groups, was calculated using the formula:

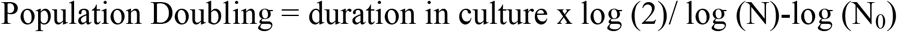

Where, N_0_ was the initial number of cells seeded, which was day 0 and N was the number of cells obtained at confluence. Cumulative PD (CPD) was compared between chondrocytes and CP from p0 to p10 (Fig 4).

**Fig 4:**
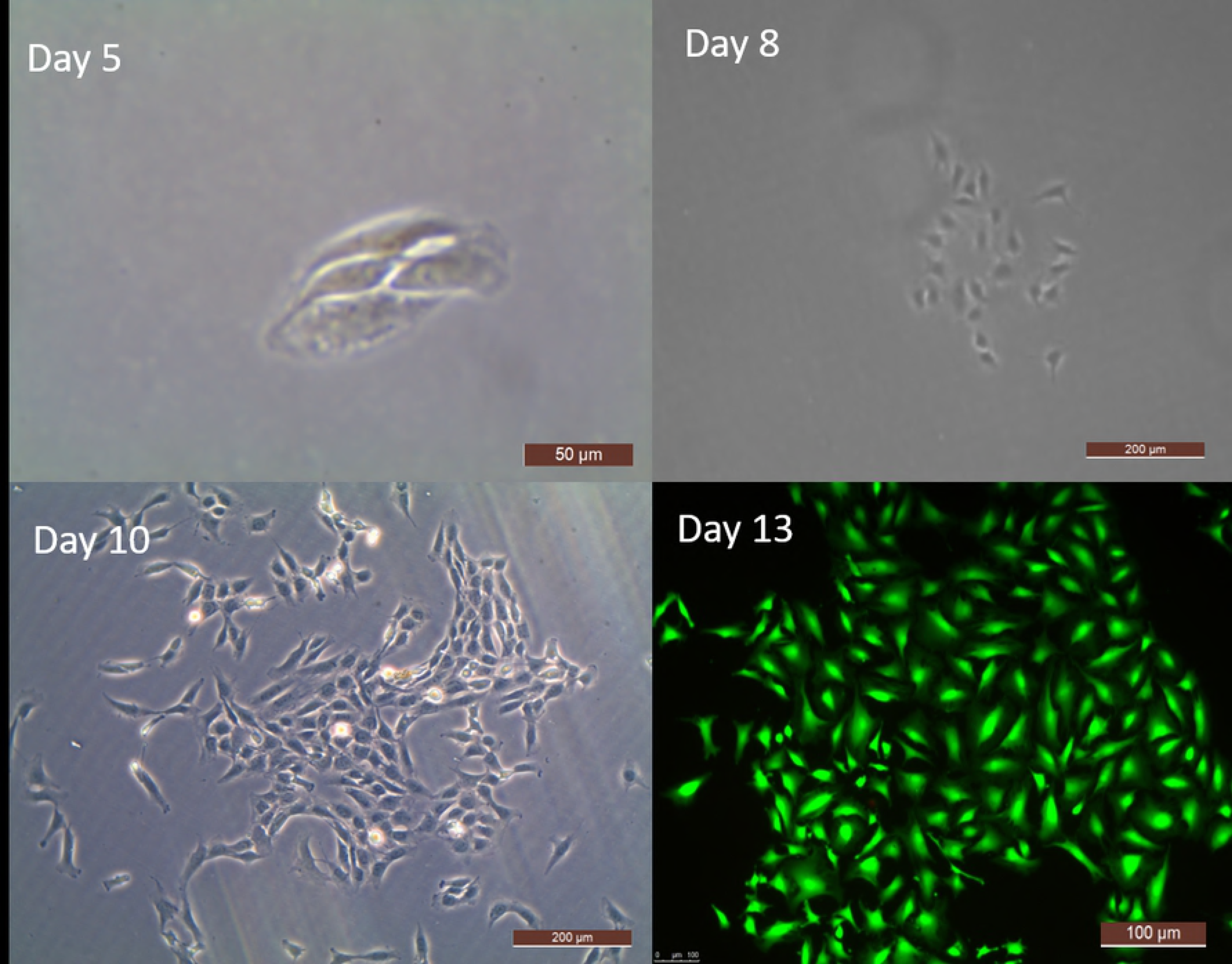
Cumulative population doubling (CPD). Data representing values for all subgroups (n=3) across time in culture (p0-p10). Data expressed as Mean±SEM. N: normal OA: osteoarthritic, C: chondrocytes, CP: chondroprogenitors and p: passage.

### Phenotyping-FACS

Chondrocytes and CPs from p0 to p10 were characterized by FACS. The studied antibodies against human surface antigen were CD105, CD73, CD90, CD34, CD45, CD14, CD54, CD44, CD9, CD106, CD29, CD151, CD49e, CD166 and CD146 (S1 Table). The staining method followed instructions provided with the manual received with individual antibodies. Harvested human chondrocytes and CPs were directly incubated with phycoerythrin (PE), fluorescein isothiocyanate (FITC) and allophycocyanin (APC) labeled antihuman antibody specific for the above-mentioned antibodies. BD FACS Calibur or BD FACS Celesta flow cytometers were used for data acquisition. Gating and compensation was applied using BD FACS Diva v 5.0.2 software and isotype controls were run for the specific CD markers. Flow cytometric analysis results are reported as Mean ± SEM (S2 and S3 Table).

### RT-PCR

p0 and p5 chondrocytes and CPs from normal and OA joints were used for RT-PCR analysis. Total RNA was isolated using TRIzol reagent (Sigma) as per manufacturer’s instructions. The nucleic acid concentration was quantified using Nanodrop 2000c spectrophotometer (ThermoScientific). Visual assessment of total RNA was done by Image Quant 400 Gel Doc system (GB) for 28s:18s ratio using 1% agarose gel containing ethidium bromide. 200 ng of RNA (in 10μl reaction volume) was reverse transcribed to cDNA using RT-RTCK-03 kit (Eurogenetec). The RT cycle conditions using Gene Amp PCR System 9700 were as follows: 25°C for 10 minutes, 48°C for 30 minutes and 95°C for 5 minutes. Quantitative RT-PCR was performed with Takyon™ Rox SYBR Master Mix dTTP Blue (Eurogenetec) by QuantStudio 6K Flex (Applied Biosystem). Cycling conditions for acquiring fluorescence were as follows: 95°C for 3 minutes Takyon™ activation, 40 amplification cycles (95°C for 3 seconds and annealing at 60°C). Threshold cycle (Ct) value was defined as the cycle number at which the curve crossed the threshold set at the mid-point of the log fluorescence expansion and each sample was run in triplicates. Relative expression for each gene was normalized to GAPDH expression. Sequences of the primers used for this study are listed in S4 Table.

### Live dead assay: Calcein AM-Ethidium homodimer

Cell viability was assessed for the four groups up to p5 on reaching a confluence of 2/3^rd^ of the flask area, using Calcein AM (for live cells) and Ethidium homodimer (for dead cells). Cells were washed with 1X PBS and incubated with 0.4μM fluorescent green Calcein AM for 30 minutes followed by Ethidium homodimer 2 μM fluorescent red for 5 minutes. Cells were subsequently washed again in PBS and observed under Leica immunofluorescence microscope (Fig 2).

### Tri-lineage Differentiation potential

Chondrocytes and CPs from normal and OA joints at p2 were assessed for multi-lineage potential. Adipogenic and osteogenic potential were evaluated in 2D cultures where cells were grown to 50% confluence prior to differentiation. Negative control which included cells cultured for the same time period with standard culture medium, were also assessed. Chondrogenic differentiation potential of all the cell groups in this study was done using three-dimensional pellet system cultures (1x 10^6^ cells). Cells were allowed to stabilize in standard growth medium for 48 hours and subjected for trilineage differentiation. Complete differentiation of human chondrocytes and CPs into a) adipocytes was performed using HiAdipoXL™ (Himedia-AL521) b) osteocytes was performed using, HiOsteoXL™ (Himedia-AL522) and c) chondrocytes was performed using HiChondroXL™ (Himedia-AL523) differentiation kits in accordance with the manufacturer’s protocol. Medium was changed once every 48-72 hours for 21 days following which differential staining was performed (Fig 5).

**Fig 5:**
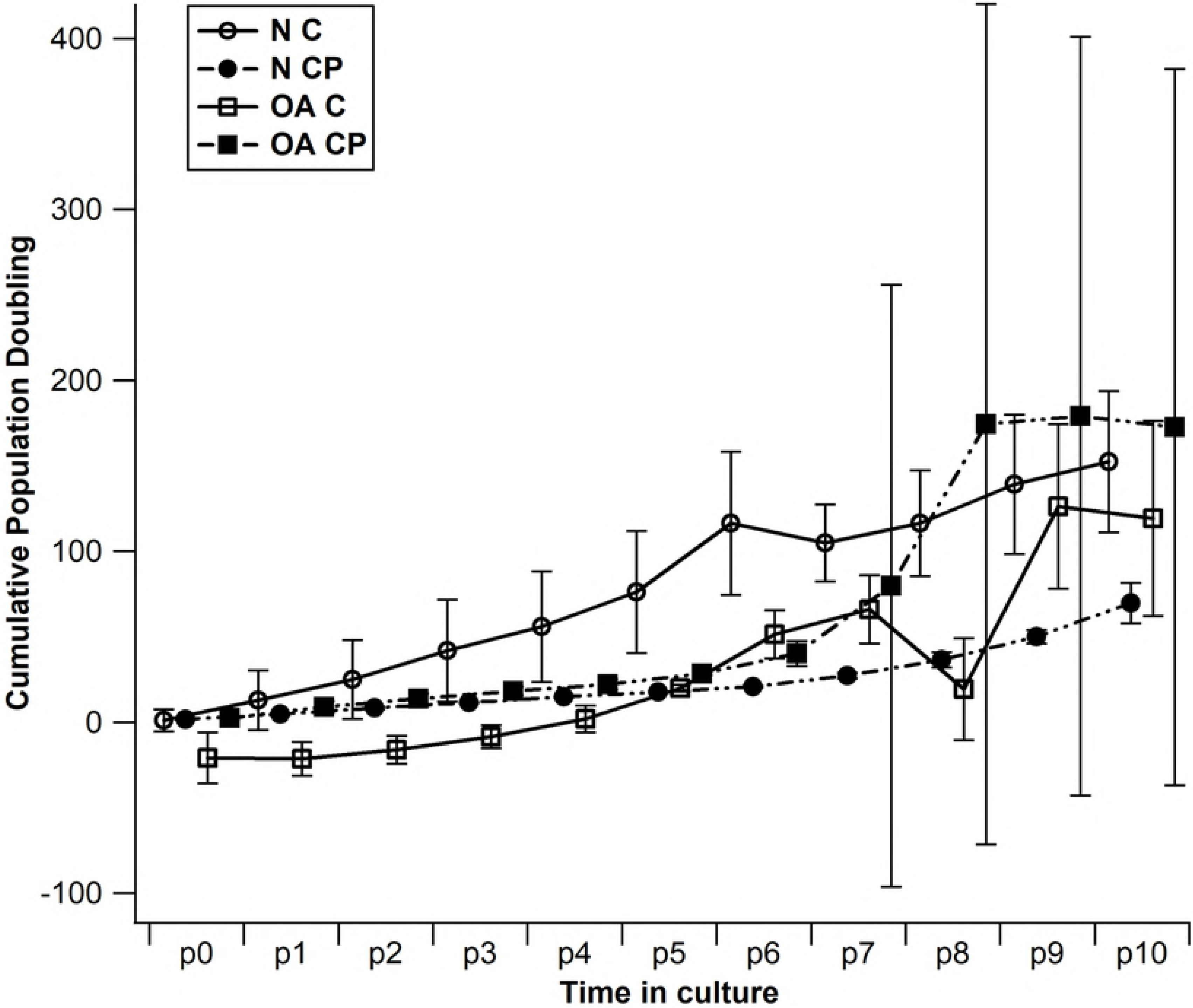
Trilineage differentiation of passage 2 chondrocytes and chondroprogenitors from normal and osteoarthritic joints. Representative microscopic images of Oil Red O (A-B) and Von Kossa (C-D) staining to confirm adipogenic and osteogenic differentiation (20X). Chondrogenic differentiation (E-F) was confirmed by Alcian Blue staining of formed cell pellets(10X).

### Differential stains- Oil Red O and Alizarin Red

The differentiated adipocytes were fixed with 10% formaldehyde for 1 hour, washed and stained with Oil Red O (Sigma) (Fig 5A-5B) and differentiated osteocytes were fixed with 70% ethanol for 1 hour, washed and stained with Alizarin Red (Sigma) (Fig 5C-5D). Images were captured using Olympus virtual slide system.

### Histological staining- Alcian Blue

To visualize glycosaminoglycan accumulation, chondrogenic pellets were fixed with 10% formaldehyde for 10 minutes, washed and stained with Alcian Blue and counterstained with neutral red (Fig 5E-5F).

### Statistical Analysis

Results following analysis of FACS, RT-PCR and CPD data were reported as mean ± standard error mean. SPSS software (version <u>22.0</u>) was used for statistical analysis. One-way ANOVA with Bonferroni correction was used to compare CPD from p0 to p10. Wilcoxon rank-sum (Mann Whitney) test was used to compare CD marker expression across different cell types (chondrocytes vs. CP) and cell source (Normal vs. OA) at p0 through p10. One-way ANOVA with Bonferroni correction was used to compare unit change of expression across all groups, using Normal CP as reference to study effect at each passage and keeping p0 as reference to study effect of time in culture. Relative expression level for each gene (normalized to the GAPDH) across different cell types (chondrocytes vs. CP) and cell source (Normal vs. OA) was compared using Wilcoxon rank-sum (Mann Whitney) test. Wilcoxon signed rank test was used to compare the relative expression level for each gene (normalized to the GAPDH) across time in culture (p0 vs. p5). A P value of < 0.05 was considered as significant.

## Results

### CPD

When CPD was compared across cell groups at each passage from p0 to p10, there was no significant difference observed in the proliferative capacity between them with all groups showing progressive increase in their growth kinetics upto p6 (Fig 4).

### FACS

FACS of both chondrocytes and CPs from normal and OA cartilage at various passages (24 hours for chondrocytes and p0, p1, p2, p3, p4, p5, p7 and p10 for both) was performed to study surface marker expression based on the categories mentioned earlier. The grading of expression utilized was <5%: nil expression, 6-35%: mild expression, 36-65%: moderate expression, 65-95%: high expression and >95% very high expression.

In the first category a) CD105: all groups showed mild to high expression except for normal C at 24hrs, p4, p10, OA chondrocytes at 24hrs and OA CP at p10 showing nil expression; b) CD73: most groups showed a very high expression with a few showing moderate to high expression; c) CD90: all groups showed a very high expression except for chondrocytes subgroups at 24 hrs which showed mild expression; d) CD106: all groups showed nil to mild expression except for normal C p1 and OA chondrocytes p0 which showed moderate expression; e) CD45: all groups showed nil expression; f) CD34: all CP subgroups showed nil expression and chondrocytes subgroups showed nil to mild expression; g) CD14: all CP subgroups showed nil expression and chondrocytes subgroups showed nil to mild expression. In the second category a) CD54: all groups showed high expression and b) CD44: all groups showed a very high expression except for normal chondrocytes at 24 hrs showing high expression and OA chondrocytes at 24hrs showing moderate expression. In the third category a) CD9: all groups showed high to very high expression; b) CD29: all groups showed a very high expression except for normal chondrocytes at P7 and OA chondrocytes at 24 hrs showing high expression; c) CD151: all groups showed a very high expression except for normal chondrocytes at 24 hrs showing high expression and OA chondrocytes at 24hrs showing moderate expression; d) CD49e: all groups showed high to very high expression except OA chondrocytes at 24 hrs showing only moderate expression; e) CD166: all groups showed high to very high expression except normal chondrocytes at 24hrs and P0 showing moderate expression and OA chondrocytes at 24hrs showing mild expression and f) CD146: all C subgroups showing mild expression except for OA chondrocytes P7 showing moderate expression and all CP subgroups in early passages showing mild to moderate expression with an upregulation in later passages.

### Intergroup comparison

When surface marker expression was compared across each passage there was no significant difference observed between chondrocytes and CPs derived from normal or OA cartilage except for the following: in the first category a) CD90 expression was higher in normal CP P0 than in normal C P0 (P=0.037, Fig 6C), b) CD34 expression was higher in normal chondrocytes P5 than in normal CP P5(P=0.046, Fig 7A) and c) CD45 expression was higher in normal chondrocytes P1 than in normal CP P1(P=0.046, Fig 7B). In the third category a) CD49e expression was higher in normal CP P5 than in normal chondrocytes P5 (P=0.046, Fig 8D) and b) CD166 expression was higher in normal CP than in normal chondrocytes at P1, P2 and P5 (P=0.046, P=0.046 and P=0.046 respectively, Fig 9A) and higher in OA CP than in OA chondrocytes at P1 and P2 (P=0.037 and P=0.037 respectively, Fig 9A).

**Fig 6:**
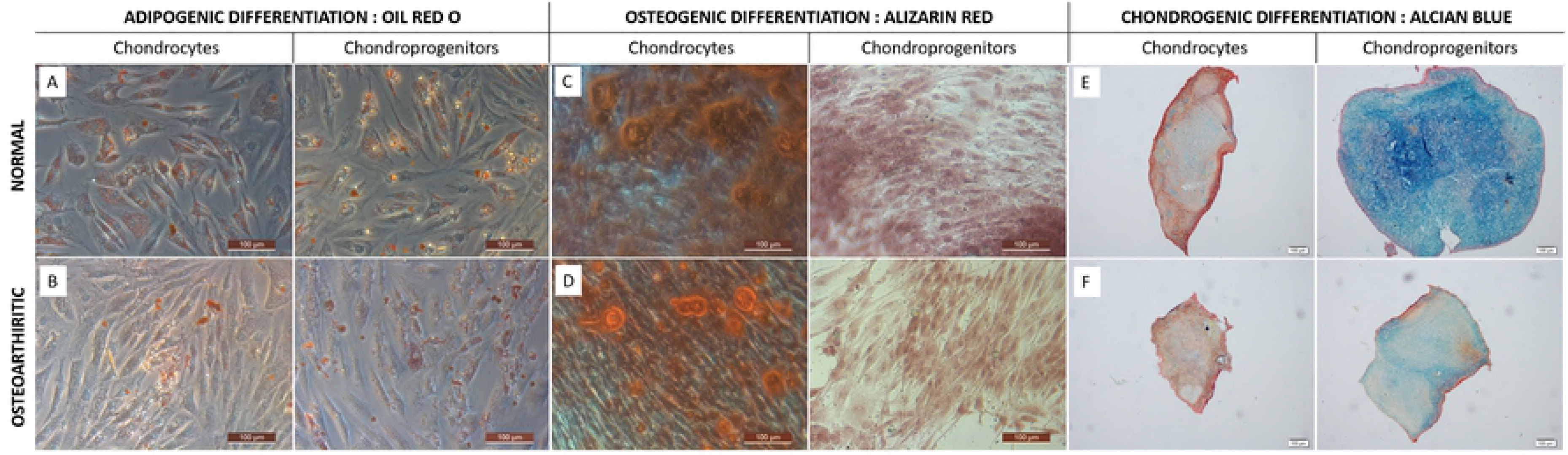
Percentage expression of CD105, CD73, CD90 and CD106 (positive MSC markers). Comparison across different cell types and cell source at p0 through p10. Data expressed as mean ± SEM (*P<0.05 using Wilcoxon rank-sum/Mann Whitney test). N: normal, OA: osteoarthritic, C: chondrocytes, CP: chondroprogenitors, p: passage.

**Fig 7:**
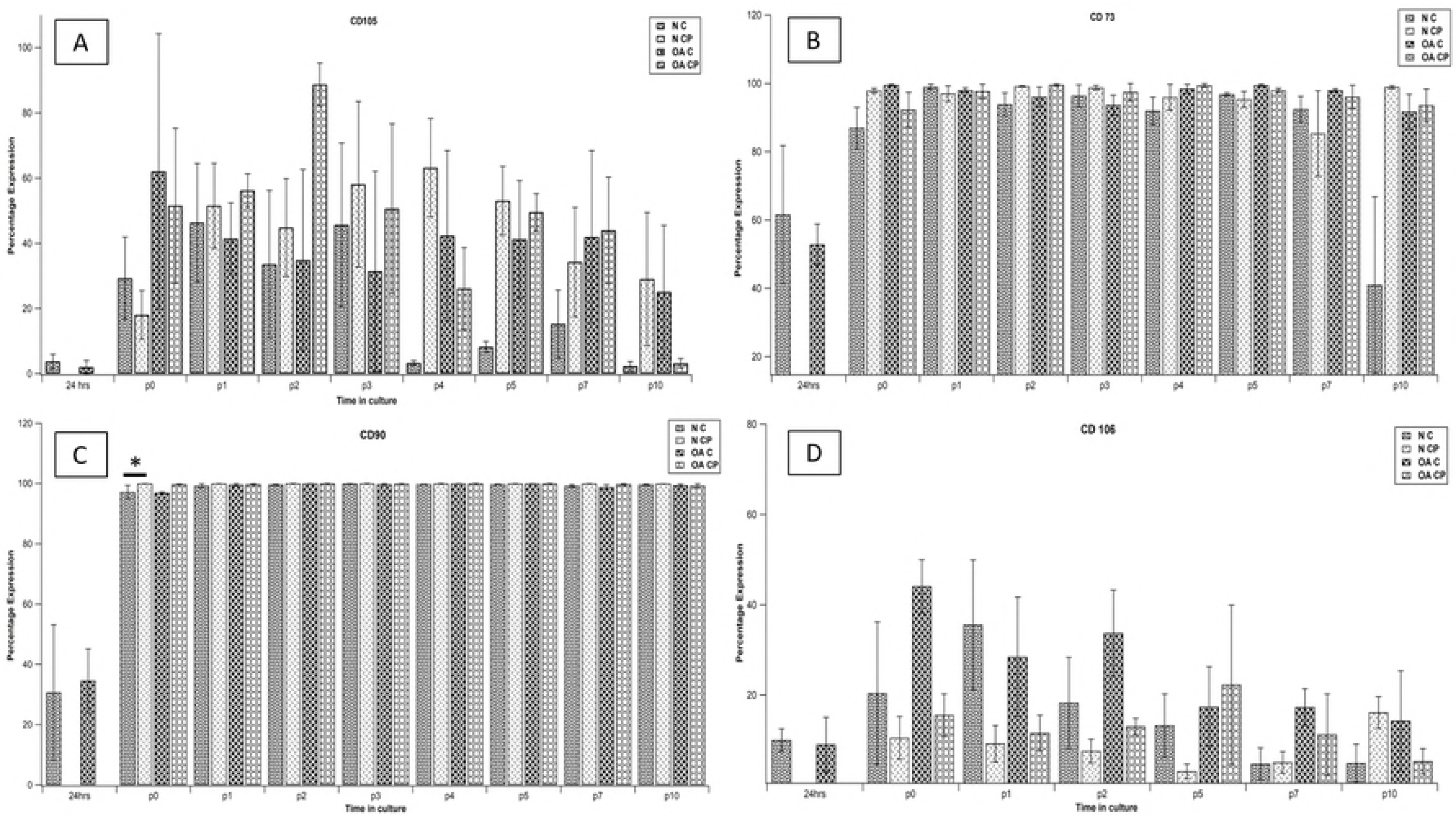
Percentage expression of CD34, CD45 and CD14 (negative MSC markers). Comparison across different cell types and cell source at p0 through p10. Data expressed as mean ± SEM (*P<0.05 using Wilcoxon rank-sum/Mann Whitney test). N: normal, OA: osteoarthritic, C: chondrocytes, CP: chondroprogenitors, p: passage.

**Fig 8:**
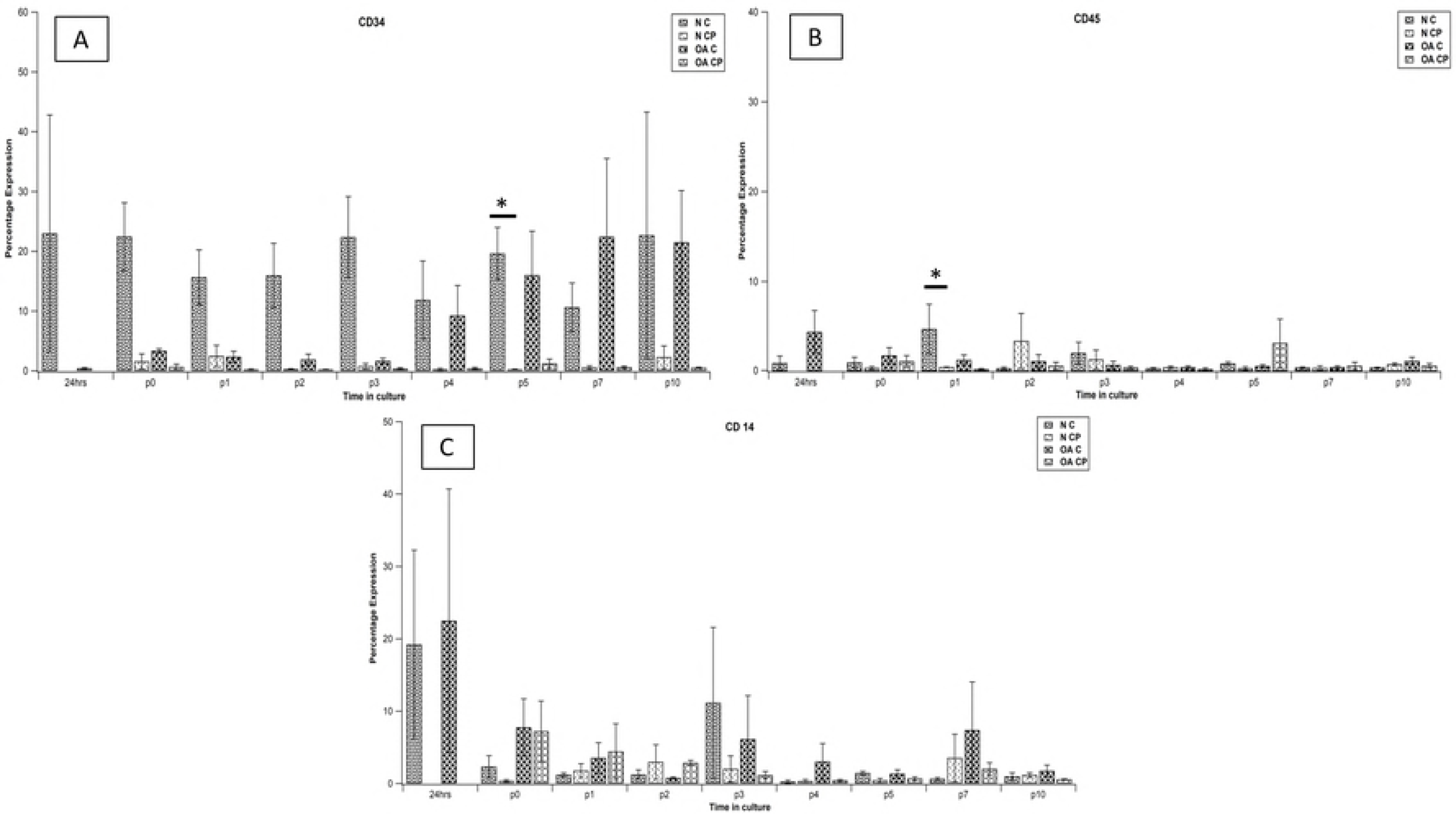
Percentage expression of CD9, CD29, CD151 and CD49e (markers of chondrogenic potential). Comparison across different cell types and cell source at p0 through p10. Data expressed as mean ± SEM (*P<0.05 using Wilcoxon rank-sum/Mann Whitney test). N: normal, OA: osteoarthritic, C: chondrocytes, CP: chondroprogenitors, p: passage.

**Fig 9:**
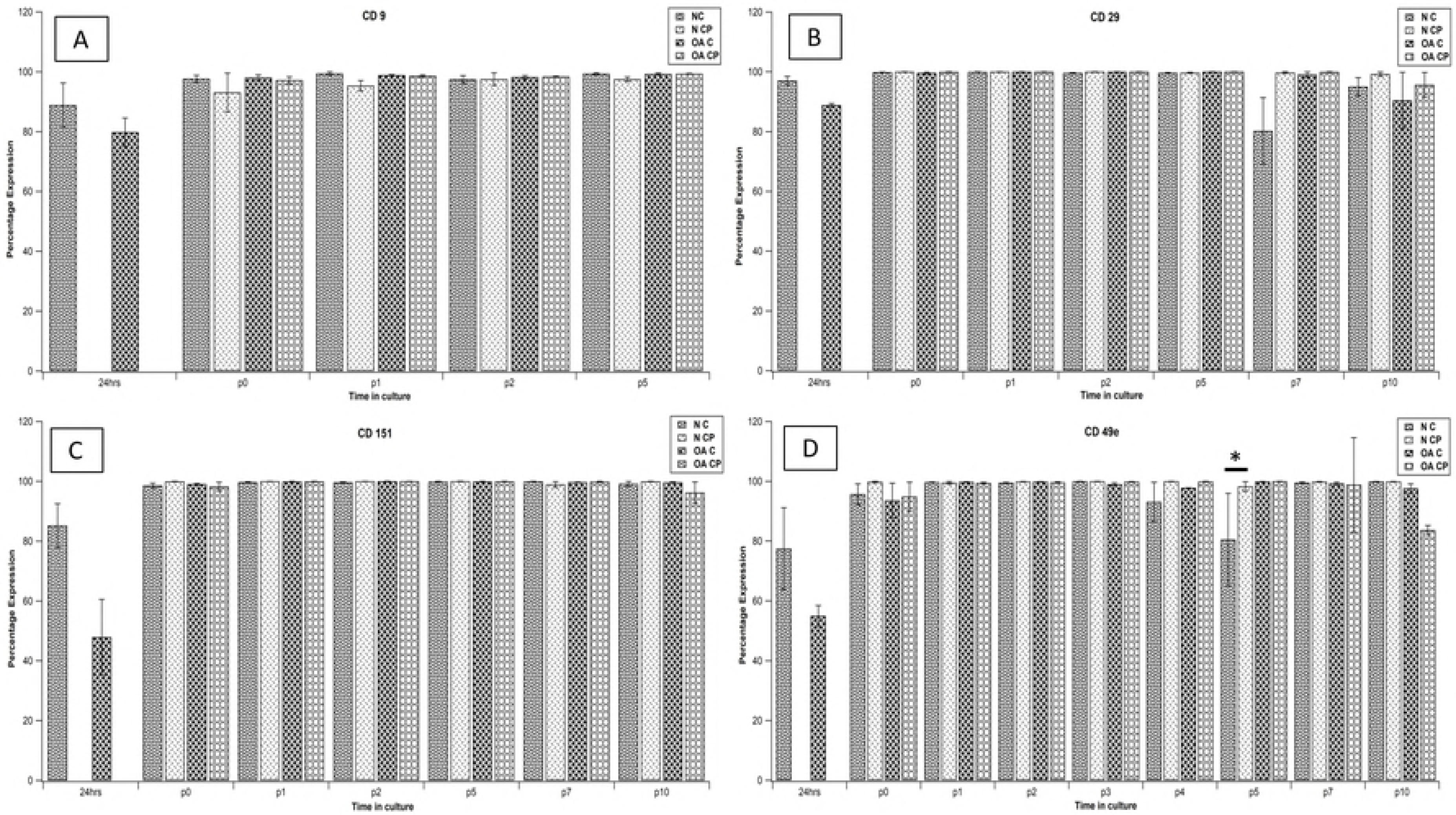
Percentage expression of CD166 and CD146 (markers of chondrogenic potential). Comparison across different cell types and cell source at p0 through p10. Data expressed as mean ± SEM (*P<0.05 using Wilcoxon rank-sum/Mann Whitney test). N: normal, OA: osteoarthritic, C: chondrocytes, CP: chondroprogenitors, p: passage.

For further analysis, normal CP values were used as reference and unit change of expression was compared across all groups at various passages. There was no significant difference observed between the groups except in CD34, CD166 and CD146. CD34 expression was higher in normal chondrocytes than in normal CP at P0, P1, P2 and P3 (P=0.006, P=0.034, P=0.022 and P=0.012 respectively, Fig 10 and Fig 11). CD166 expression was higher in normal CP than in normal chondrocytes at P0, P2 and P5(P=0.003, P=0.009 and P=0.037 respectively, Fig 11). CD146 expression was higher in normal CP than in normal chondrocytes at P1 and P7 (P=0.013 and P=0.003 respectively, Fig 11). Similarly, the expression was higher in normal CP than OA C at P1, P2 and P5 (P=0.004, P=0.023 and P=0.017 respectively, Fig 11) and OA CP at P1(P=0.002, Fig 11)

**Fig 10:**
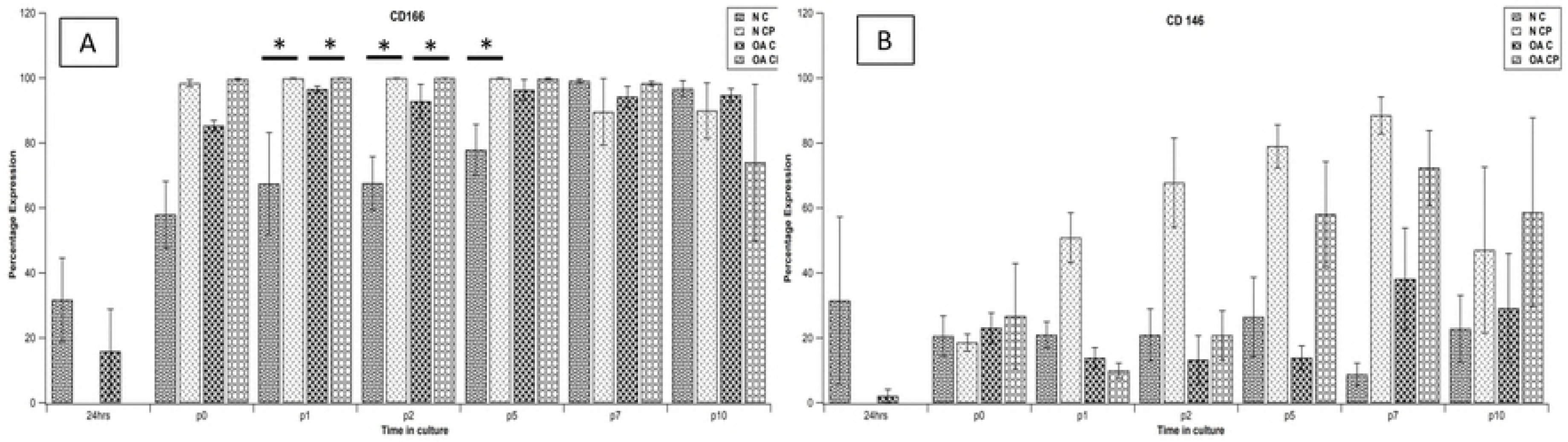
Flow cytometry values for positive, negative and chondrocyte markers, expressed as unit change across all groups, using N CP values as reference to study effect at each passage. Data expressed as mean ± SEM (Highlighted values: P<0.05 using One-way ANOVA with Bonferroni correction). N: normal, OA: osteoarthritic, C: chondrocytes, CP: chondroprogenitors, p: passage.

**Fig 11:**
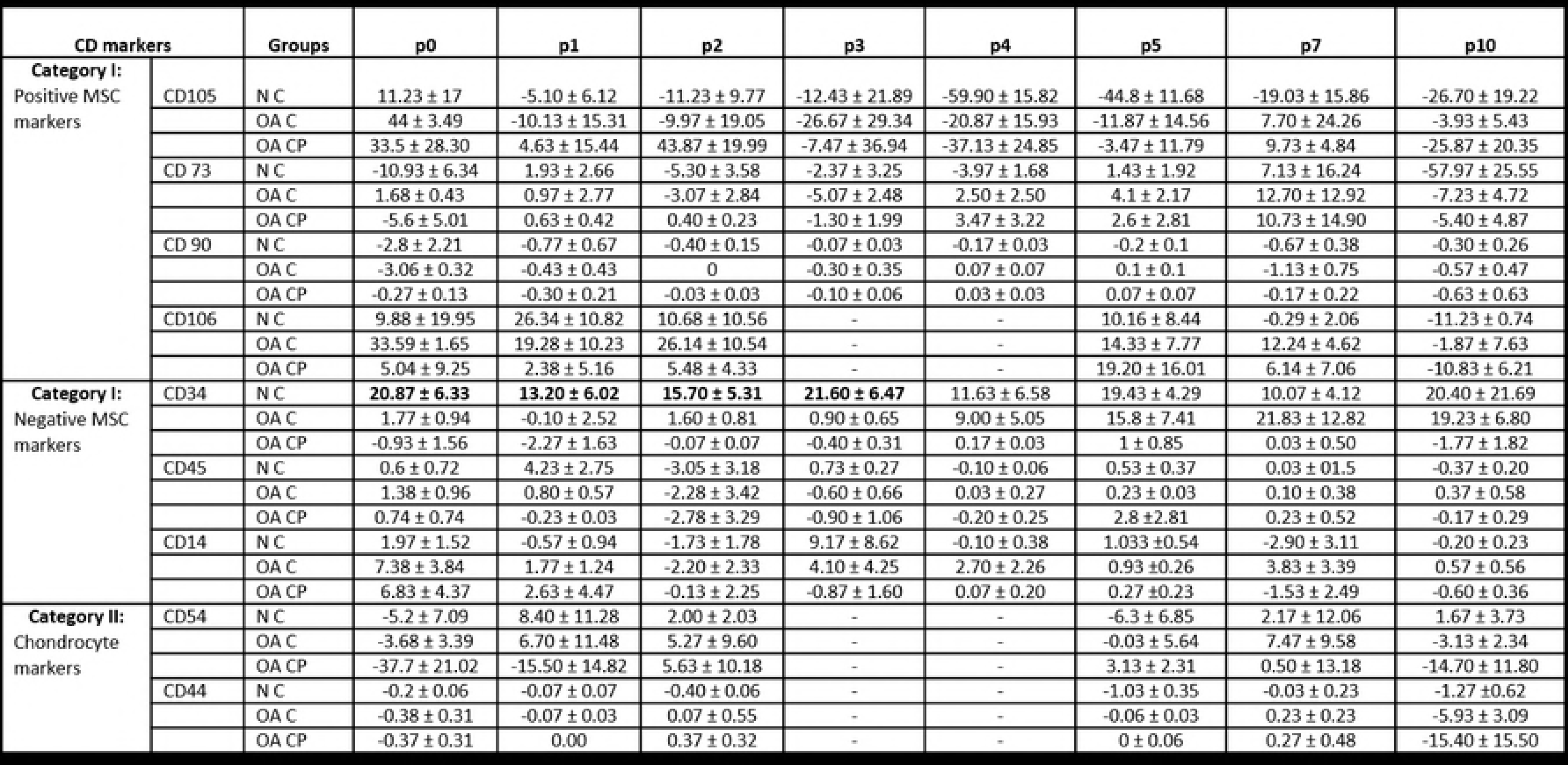
Flow cytometry values for markers of enhanced chondrogenesis, expressed as unit change across all groups, using N CP values as reference to study effect at each passage. Data expressed as mean ± SEM (Highlighted values: P<0.05 using One-way ANOVA with Bonferroni correction). N: normal, OA: osteoarthritic, C: chondrocytes, CP: chondroprogenitors, p: passage.

To study the effect of time in culture on surface marker expression across the four groups, P0 values of CD markers for each group were used as reference and unit change of expression was compared for each CD marker within the group. There was no significant difference observed across passages in the groups except in a) normal chondrocytes, CD90 expression was higher at P0 than at 24hours (P=0.000, Fig 12), b) Normal CP, CD146 expression was higher in P7 than at P0 (P=0.034, Fig 13) and c) OA chondrocytes; CD73, CD90, CD44, CD151, CD9, CD166 and CD49e expressions were higher at P0 than at 24hours (P=0.000, P=0.000, P=0.001, P=0.000, P=0.001, P=0.000 and P=0.000 respectively, Fig 12 and Fig 13)

**Fig 12:**
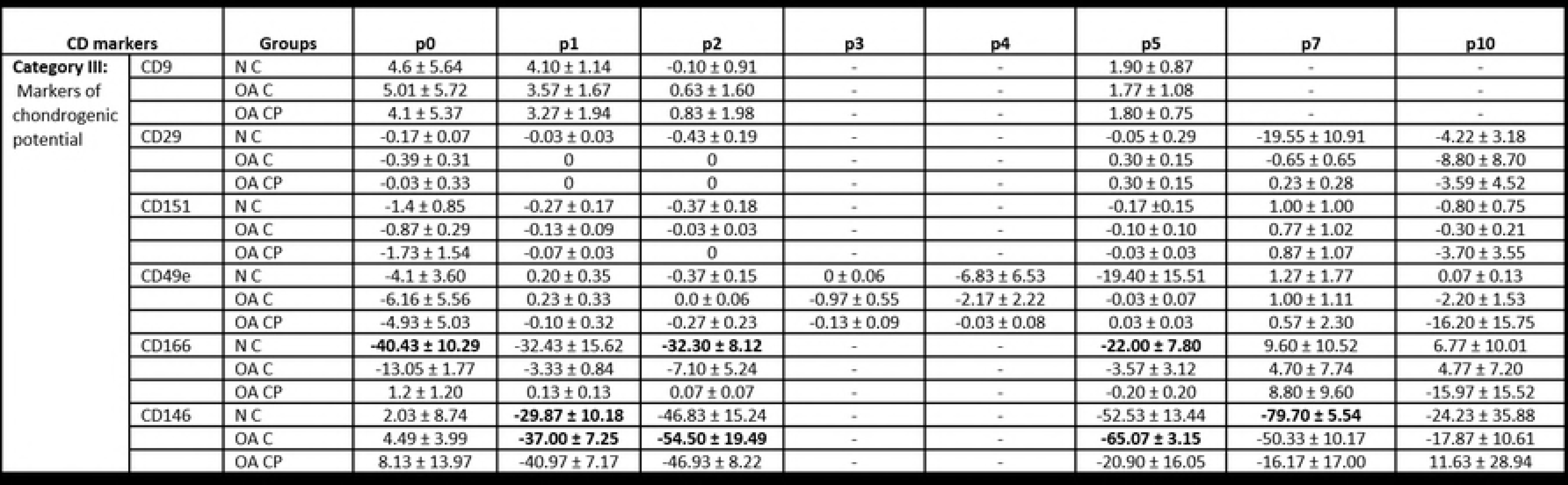
Flow cytometry values for positive, negative and chondrocyte markers, expressed as unit change across all groups using p0 values as reference to study effect of time in culture. Data expressed as mean ± SEM (Highlighted values: P<0.05 using One-way ANOVA with Bonferroni correction). N: normal, OA: osteoarthritic, C: chondrocytes, CP: chondroprogenitors, p: passage.

**Fig 13:**
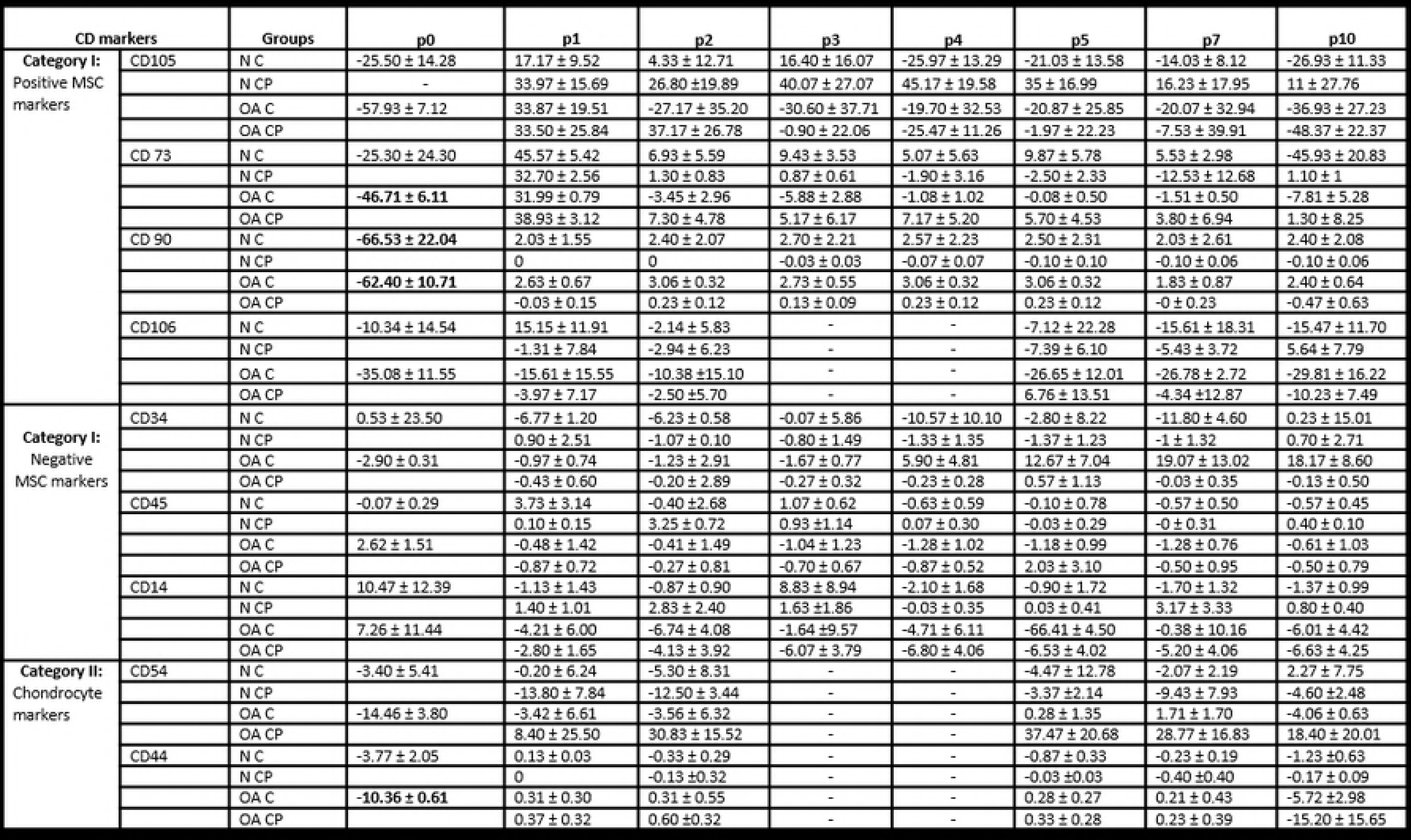
Flow cytometry values for markers of enhanced chondrogenesis, expressed as unit change across all groups using p0 values as reference to study effect of time in culture. Data expressed as mean ± SEM (Highlighted values: P<0.05 using One-way ANOVA with Bonferroni correction). N: normal, OA: osteoarthritic, C: chondrocytes, CP: chondroprogenitors, p: passage.

### mRNA expression of chondrogenic and hypertrophy markers

Gene expression of specific primers in both chondrocytes and CPs from normal and OA cartilage at P0 and P5 were examined to study the levels of chondrogenic and hypertrophy markers. RT-PCR analysis of chondrogenic markers showed that both cell populations demonstrated a high expression of Aggrecan and moderate to high expression of SOX9. Collagen II expression was high in two chondrocytes subgroups (normal chondrocytes P0 and OA chondrocytes P0), moderate in two CP subgroups (normal CP P0 and OA CP P0) and undetermined in all other subgroups. Analysis of hypertrophy markers show that both cell populations demonstrated a high expression of Collagen I, moderate to low expression of MMP-13 and low expression of RUNX2. Collagen X expression was seen to be low in the subgroup (OA C P0) and undetermined in all others. When relative gene expression was compared across P0 and P5 there was no significant difference observed between chondrocytes and CPs derived from normal or OA cartilage except for in a) Collagen II OA chondrocytes P0(Mean ΔCt: 6.01 ± 1.76) demonstrated a higher expression than OA CP P0 (Mean ΔCt: 14.84 ± 0.69, P=0.049, Fig 14),b) Aggrecan normal chondrocytes P5 (Mean ΔCt: 1.53 ± 0.086) demonstrated a higher expression than normal CP P5 (Mean ΔCt: 5.97 ± 0.74, P=0.049, Fig 15),c) SOX9 OA CP P5 (Mean ΔCt: 7.81 ± 0.20) demonstrated a higher expression than normal CP P5 (Mean ΔCt: 8.56 ± 0.12, P=0.049, Fig 15),d) Collagen X OA chondrocytes P0 (Mean ΔCt: 16.15 ± 0.89) demonstrated a higher expression than OA CP P0 (Mean ΔCt: 20.34 ± 1.13, P=0.049, Fig 14) and e) RUNX2 OA C P0 (Mean ΔCt: 14.22 ± 0.65) demonstrated a higher expression than normal chondrocytes P0 (Mean ΔCt: 17.11 ± 0.64, P=0.049, Fig 14). When mRNA expression was compared to study the effect of time in culture for both cell types, no significant difference was observed in the relative expression in the various chondrocyte and CP sub-groups (Fig 16 and 17)

**Fig 14:**
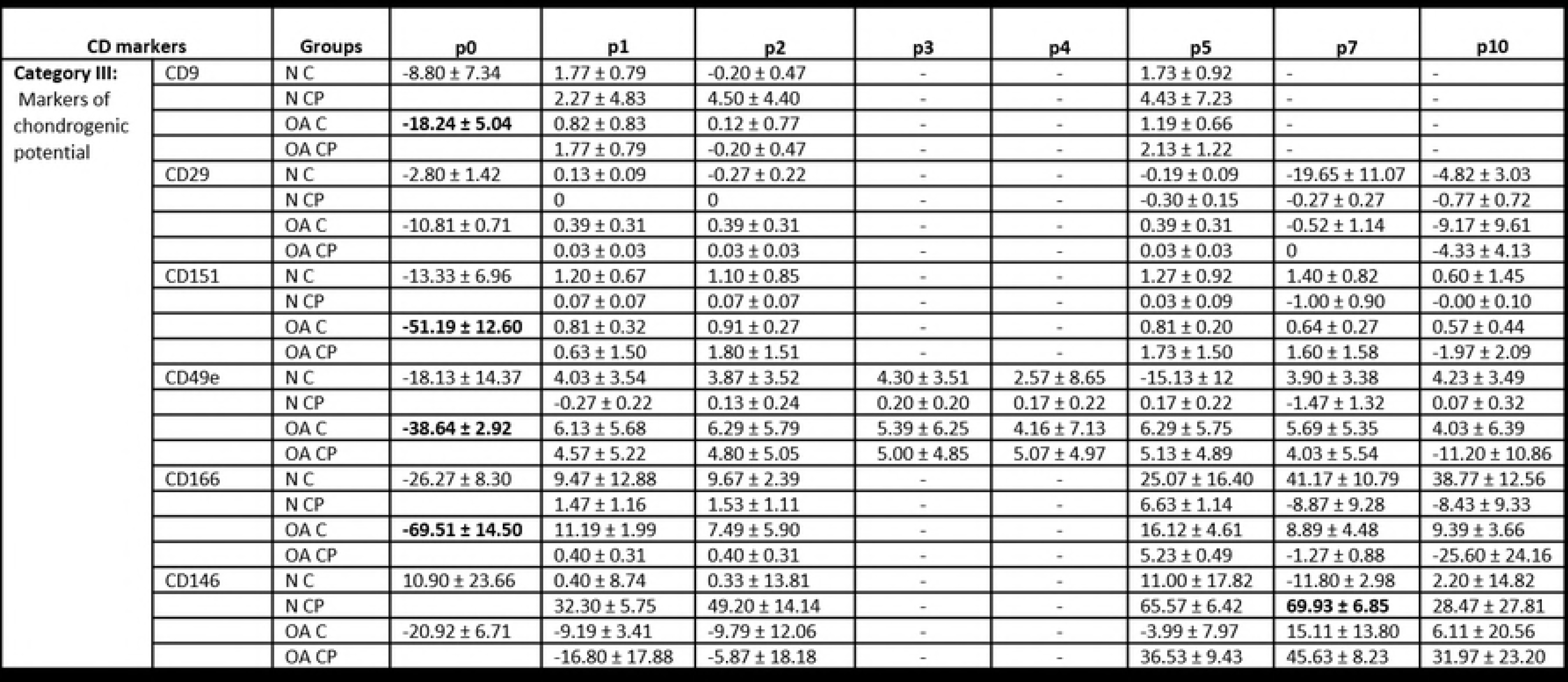
Relative expression of Collagen II, Aggrecan, SOX9, Collagen I, Collagen X, RUNX2 and MMP-13 in all subgroups at passage 0. Level for each gene across different cell types (C vs. CP) and cell source (N vs. OA) was compared using Wilcoxon rank-sum (Mann Whitney) test. ΔCt values normalized to GAPDH are expressed as Mean ± SEM (*P<0.05). N: normal, OA: osteoarthritic, C: chondrocytes, CP: chondroprogenitors, p0: passage 0 and p5: passage 5. Samples taken from n=3 donors, each sample(n) was run in triplicates.

**Fig 15:**
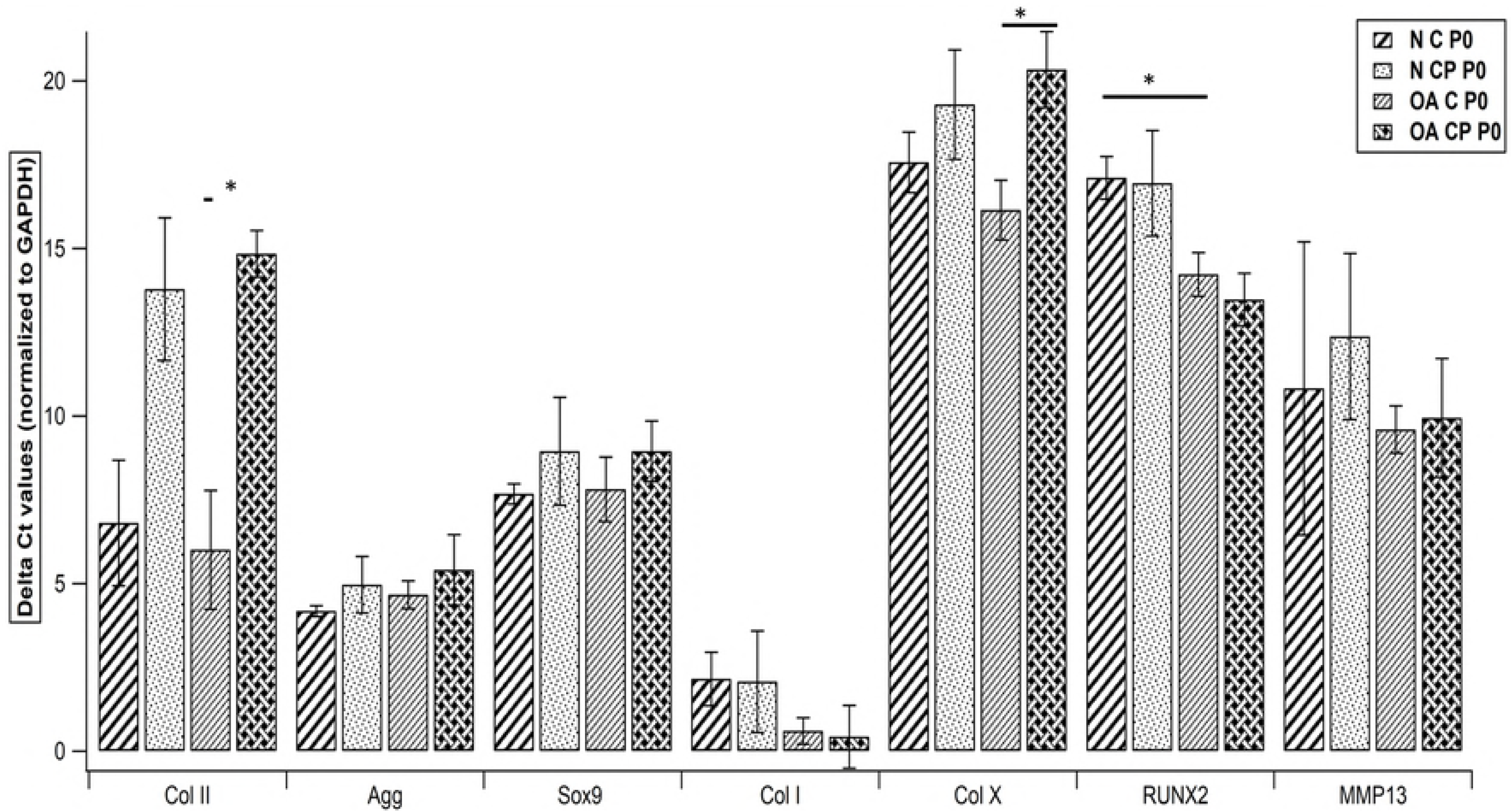
Relative expression of Collagen II, Aggrecan, SOX9, Collagen I, Collagen X, RUNX2 and MMP-13 in all subgroups at passage 0. Level for each gene across different cell types (C vs. CP) and cell source (N vs. OA) was compared using Wilcoxon rank-sum (Mann Whitney) test. ΔCt values normalized to GAPDH are expressed as Mean ± SEM (*P<0.05). N: normal, OA: osteoarthritic, C: chondrocytes, CP: chondroprogenitors, p0: passage 0 and p5: passage 5. Samples taken from n=3 donors, each sample(n) was run in triplicates.

**Fig 16:**
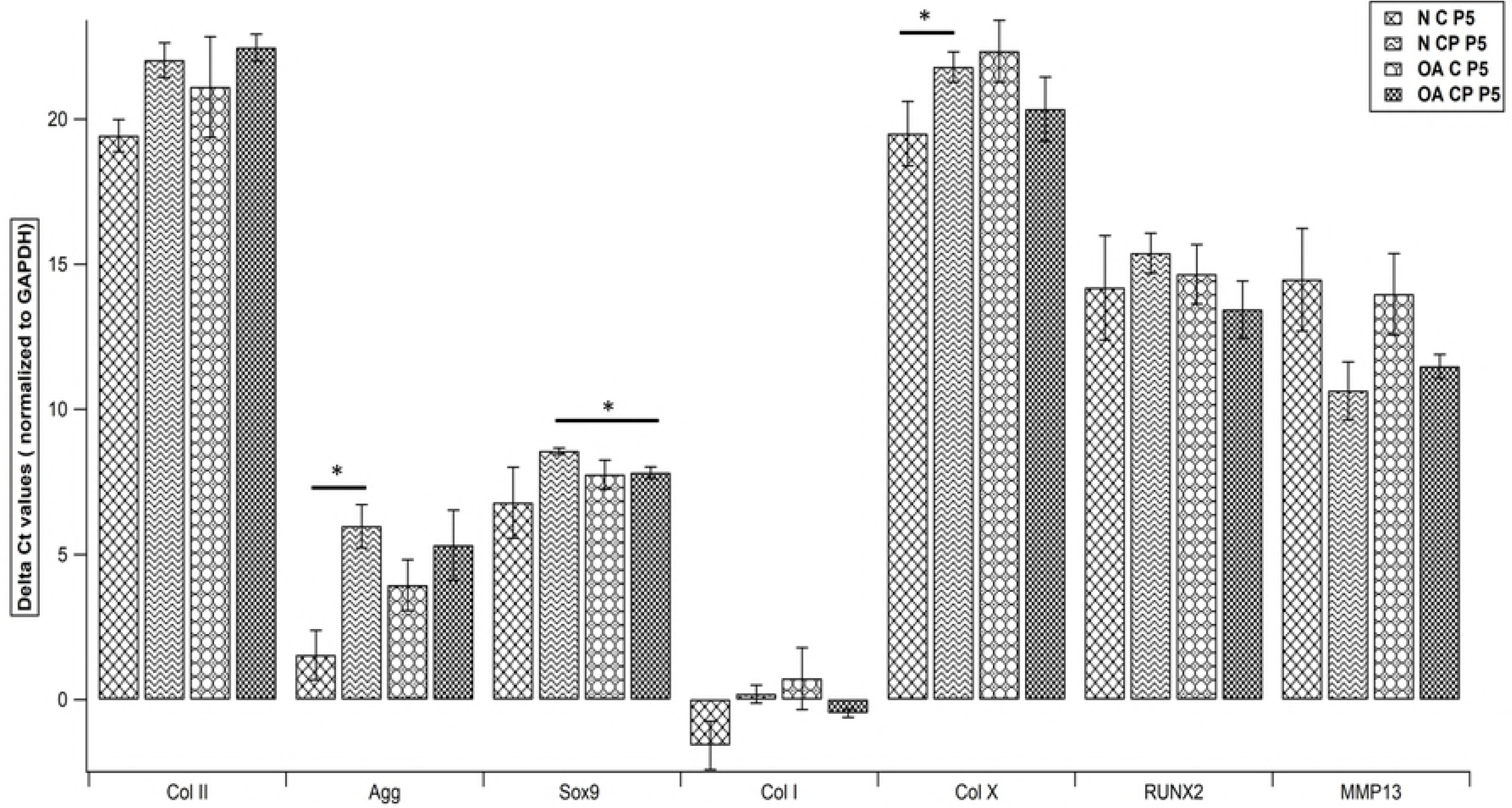
Relative expression of Collagen II, Aggrecan, SOX9, Collagen I, Collagen X, RUNX2 and MMP-13 in all chondrocyte subgroups across time in culture (p0 vs. p5). Data expressed as Mean±SEM (Wilcoxon sign rank test). N: normal, OA: osteoarthritic, C: chondrocytes, p0: passage 0, p5: passage 5. Samples taken from n=3 donors, each sample(n) was run in triplicates.

**Fig 17:**
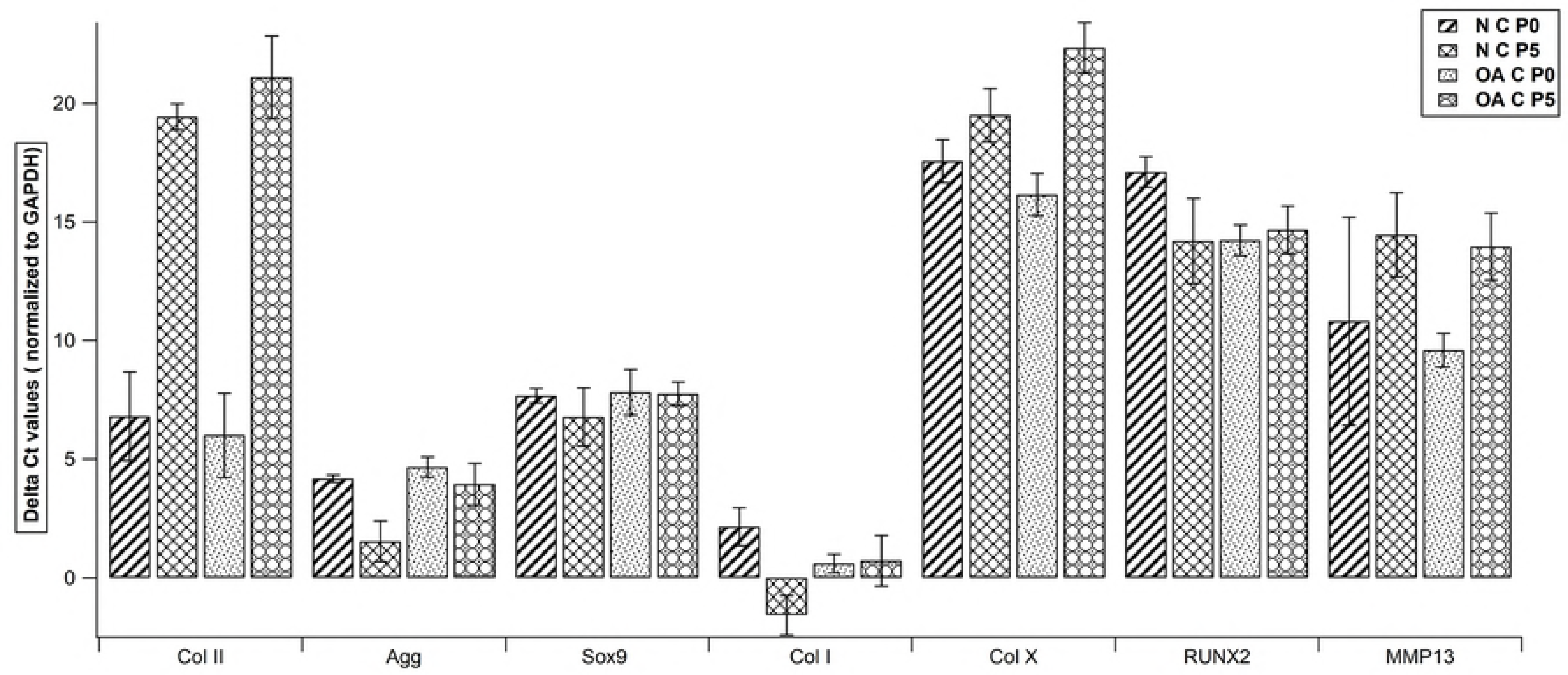
Relative expression of Collagen II, Aggrecan, SOX9, Collagen I, Collagen X, RUNX2 and MMP-13 in all chondroprogenitor subgroups across time in culture (p0 vs. P5). Data expressed as Mean±SEM (Wilcoxon sign rank test). N: normal, OA: osteoarthritic, C: chondrocytes, p0: passage 0, p5: passage 5. Samples taken from n=3 donors, each sample(n) was run in triplicates.

### Tri-lineage Differentiation

Both chondrocytes and CPs derived from normal and OA cartilage displayed tri-lineage differentiation potential. Qualitative analysis (Oil red O for lipid vacuole accumulation) showed no difference between the two populations when adipogenic potential was assessed (Fig 5A-B). Alizarin red staining indicative of mineralization used to assess osteogenic potential showed positive staining in both cell groups with higher uptake seen with chondrocytes (Fig 5C-D). Alcian blue staining, suggestive of glycosaminoglycan deposition used for assessing chondrogenic differentiation showed positive staining in both cell groups with better deposition seen with CPs. Chondrocytes and CPs derived from normal cartilage demonstrated better staining with Alcian blue when compared to their counterparts derived from OA cartilage (Fig 5E-F).

## Discussion

Articular cartilage can be regenerated from its two native cell types (chondrocytes and CPs), if their use in cell-based therapy is optimized. Extensive work on chondrocytes has afforded valuable information to their use in cartilage repair, although questions pertaining to their behavior in culture remain unanswered(13–16). On the other hand, CPs, relatively recent in the field of cell-based repair have been hailed as a promising cell type due to their proposed MSC-like characteristics and enhanced chondrogenesis. Although single source derivation of both cell populations is advantageous, lack of a characteristic differentiating marker obscures clear identification of either cell type. The importance of differentiation between the two cell types is not only necessary to create a biological profile but is also required to assess cell type superiority for cartilage regeneration.

Since CPs have been likened to MSCs(17) and chondrocytes(35) are regarded to be a mature cell-type, we first considered differentiating the two cell populations based on expression of classical MSC markers. Our results showed that both chondrocytes and CPs expressed high levels of CD73 and CD90 but varying levels for CD105 and CD106, all known positive markers (Figure 6). CD105, in addition to being an MSC marker has also been reported to be expressed in cells exhibiting higher chondrogenic potential (35–37). However, our findings suggest otherwise as CD105 expression in both cell populations was not very high with no significant difference between them. This corroborated with an earlier report demonstrating low CD105 expression in bone marrow-MSCs and human articular chondrocytes (HAC) questioning its suitability as an identifying marker of enhanced chondrogenesis(36). Regarding CD106 which has also been reported to be highly expressed in HAC, we found that both cell populations show just minimal expression albeit higher in chondrocytes though not to a significant level (Fig 6D). When expression of negative MSC markers was assessed, we found that both cell population showed low expression of CD45 and CD14 (Fig 7). However, there was a significant difference in the expression of CD34 as levels in chondrocyte groups were much higher than CP groups (Fig 7A; Table 1). This leads us to suggest that CD34, a hematopoietic stem and progenitor cell marker(37) although not expected to be expressed by cells derived from an avascular tissue like articular cartilage, may be used to distinguish chondrocytes from CPs.

The second category considered was markers (CD54 and CD44) which specifically identify chondrocytes since they are receptors for hyaluronan and have been reported to modulate chondrocyte metabolism(27,38). A steady decline in CD54/CD44 ratio has been reported in HAC when expanded in culture even during early passages(16). However, we found that both cell populations expressed high levels of CD54 and CD44 with no downregulation in expression seen with prolonged expansion in culture even upto P10(Fig 18).

**Fig 18:**
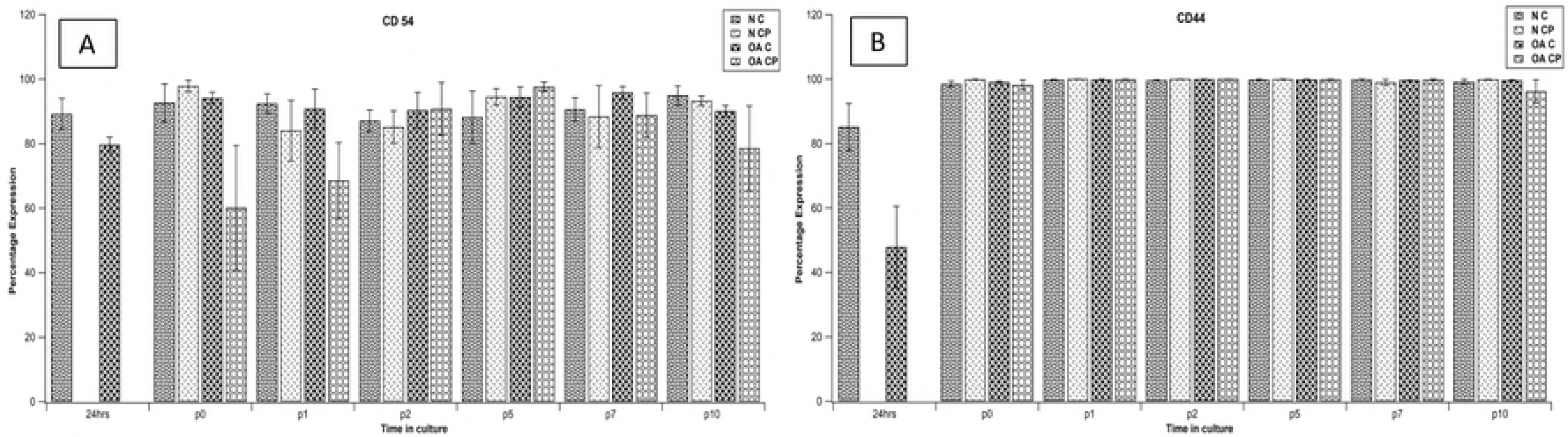
Percentage expression of CD54 and CD44 (chondrocyte markers). Comparison across different cell types and cell source at p0 through p10. Data expressed as mean ± SEM (*P<0.05 using Wilcoxon rank-sum/Mann Whitney test). N: normal, OA: osteoarthritic, C: chondrocytes, CP: chondroprogenitors, p: passage.

When expression of markers indicating chondrogenic potential was assessed we found that both cell populations exhibited high levels of CD9, CD29, CD151 and CD49e with no significant difference between the two (Fig 8). CD49e which forms a heterodimeric fibronectin receptor along with integrin β1(CD29), has been defined as a definitive marker for CPs, and forms the basis for their isolation employing fibronectin differential adhesion assay(17,28). A very important observation was that all chondrocyte groups including freshly isolated cells expressed high levels of CD49e and CD29, matching those expressed in CPs. These results indicate that CD49e is not a cell specific marker when differentiation between chondrocytes and CPs is required as has already been evidenced by earlier reports(15,39). CD166 in addition to being considered a marker of strong chondrogenic potential has also been reported to be an identifier for mesenchymal progenitor cells within human cartilage(31,40). Our results showed that even chondrocytes along with CPs expressed high levels of CD166, although expression in CPs was significantly higher than that in chondrocytes at early passages (Fig 9; Table 1). CD146, another bio-marker for enhanced chondrogenesis and reported to be expressed by early mesenchymal lineage stem cells was seen to be minimally expressed by chondrocytes(41–43). CPs on the other hand demonstrated a significantly higher expression of CD146 than chondrocytes with an upregulation in levels seen with increased time in culture (Fig 9, Table 1 and 2). From these results it may be proposed that if cells are chosen from early passages then CD166 and CD146 may be considered as positive bio-markers suitable for cell specific sorting while selecting chondroprogenitors.

Analysis of mRNA expression to assess variable difference in chondrogenic potential and degree of hypertrophy between the two cell populations yielded mixed results. While considering markers for chondrogenesis, both populations showed high levels of SOX9 and Aggrecan whereas chondrocytes from an earlier passage showed significantly higher levels of Collagen II than CPs. Evaluation of markers indicating tendency for hypertrophy revealed that both cell populations showed high expression of collagen I, moderate to low expression of MMP-13 and low expression of Collagen X and RUNX2 while CPs demonstrated a significantly lower expression of Collagen X as compared to chondrocytes. These results lend support to earlier reports which show a downregulation of Collagen II and an upregulation of Collagen I in chondrocytes when expanded in culture and additionally suggest a similar trend for CPs (38,44). Our observations are also in accord with an earlier study which suggests that CPs are a suitable contender for cell-based repair due to their lower propensity for hypertrophy(44).

In order to assess biological characteristics like replicative and differentiation potential which have been reported to be superior in CPs(17) given their MSC like nature when compared to a mature cell like chondrocyte, we compared their growth kinetics based on CPD and tri-lineage differentiation potential. The results did not show a significant difference in the proliferative capacity between the two cell populations when CPD was considered even though CPs additionally received TGFβ2 and FGF2 (growth factors) supplementation while in culture(17,18). When results for trilineage differentiation were analyzed, an interesting finding was that in addition to CPs, chondrocytes expanded in culture (p2) also demonstrated potential for adipogenic, osteogenic and chondrogenic differentiation. Qualitative comparison revealed that CPs demonstrated higher uptake of Alcian blue and lower uptake of Alizarin red as compared to chondrocytes suggesting their greater potential for chondrogenesis and lower tendency for hypertrophy.

This study was the first attempt where characterization was performed on two cell populations, not only derived from a single source but also the first to utilize human articular cartilage. A small sample size and donor to donor variability were challenges encountered and may have contributed to certain outcomes remaining obscure. The reported data also constructs a better biological profile of human CPs as well as chondrocytes since cells obtained from normal/OA joints and expanded in culture (up to p10) were used for acquiring information pertaining to parameters including FACS, gene expression, replicative potential and tri-lineage differentiation.

We observe that both cell populations exhibit similar characteristics and since CD49e does not seem to be a discrete marker of CP isolation, need for better suited bio-markers of differentiation is warranted. Our findings suggest that cell sorting based on using CD34(-), CD166(+) and CD146(+) will yield a population of cells primarily composed of CPs. This would be beneficial as the results using RT-PCR and differentiation studies also indicate that CPs demonstrated a lesser susceptibility for hypertrophy and a higher potential for chondrogenesis.

In conclusion, the study implies that CPs derived preferably from normal as opposed to OA joints and isolated using markers with higher specificity would yield translatable results in terms of enhanced chondrogenesis and reduced hypertrophy, both indispensable for the field of cartilage regeneration.

## Acknowledgment

The authors would like to acknowledge the Center for Stem Cell Research, Christian Medical College, Vellore for infrastructural support, Dr. George Thomas for critical revision of the manuscript and Mr. Bijesh Kumar Yadav (Department of Biostatistics, CMC Vellore) for help with statistical analysis.

## Supporting Information

**S1 Table: List of antibodies used for characterization of chondrocytes and chondroprogenitors by flow cytometric analysis.** MSC: mesenchymal stem cell, CD: cluster of differentiation, FITC: fluorescein isothiocyanate, PE: phycoerythrin and APC: allophycocyanin.

**S2 Table: Flow cytometric data (percentage expression) for positive, negative and chondrocyte markers, of all CD markers from various cell groups at all passages.** Data is expressed as percentage mean ± SEM (n=3).

**S3 Table: Flow cytometric data (percentage expression) for markers of enhanced chondrogenesis, of all CD markers from various cell groups at all passages.** Data is expressed as percentage mean ± SEM (n=3).

**S4 Table: Sequence of the primers used for RT-PCR**. SOX9(SRY-box9), RUNX2(Runt related transcription factor 2), MMP-13(metalloproteinase-13) and GAPDH (glyceraldehyde 3-phosphate dehydrogenase).

## References

1. Madeira C, Santhagunam A, Salgueiro JB, Cabral JMS. Advanced cell therapies for articular cartilage regeneration. Trends Biotechnol. 2015 Jan 1;33(1):35–42.

2. Grassel S, Anders S. [Cell-based therapy options for osteochondral defects. Autologous mesenchymal stem cells compared to autologous chondrocytes]. Orthopade. 2012 May;41(5):415–428; quiz 429-430.

3. Mobasheri A, Kalamegam G, Musumeci G, Batt ME. Chondrocyte and mesenchymal stem cell-based therapies for cartilage repair in osteoarthritis and related orthopaedic conditions. Maturitas. 2014 Jul;78(3):188–98.

4. Oreffo ROC, Cooper C, Mason C, Clements M. Mesenchymal stem cells: lineage, plasticity, and skeletal therapeutic potential. Stem Cell Rev. 2005;1(2):169–78.

5. Akkiraju H, Nohe A. Role of Chondrocytes in Cartilage Formation, Progression of Osteoarthritis and Cartilage Regeneration. J Dev Biol. 2015 Dec;3(4):177–92.

6. Fernandez-Moure JS, Corradetti B, Chan P, Van Eps JL, Janecek T, Rameshwar P, et al. Enhanced osteogenic potential of mesenchymal stem cells from cortical bone: a comparative analysis. Stem Cell Res Ther. 2015 Oct 26;6:203.

7. Liu X, Kumagai G, Wada K, Tanaka T, Asari T, Oishi K, et al. High Osteogenic Potential of Adipose- and Muscle-derived Mesenchymal Stem Cells in Spinal-Ossification Model Mice. Spine. 2017 Dec 1;42(23):E1342–9.

8. Akgun I, Unlu MC, Erdal OA, Ogut T, Erturk M, Ovali E, et al. Matrix-induced autologous mesenchymal stem cell implantation versus matrix-induced autologous chondrocyte implantation in the treatment of chondral defects of the knee: a 2-year randomized study. Arch Orthop Trauma Surg. 2015 Feb 1;135(2):251–63.

9. Freitag J, Li D, Wickham J, Shah K, Tenen A. Effect of autologous adipose-derived mesenchymal stem cell therapy in the treatment of a post-traumatic chondral defect of the knee. BMJ Case Rep. 2017 Oct 15;2017:bcr-2017–220852.

10. Kamel NS, Arafa MM, Nadim A, Amer H, Amin IR, Samir N, et al. Effect of intra-articular injection of mesenchymal stem cells in cartilage repair in experimental animals. Egypt Rheumatol. 2014 Oct 1;36(4):179–86.

11. Harris JD, Siston RA, Brophy RH, Lattermann C, Carey JL, Flanigan DC. Failures, reoperations, and complications after autologous chondrocyte implantation - a systematic review. Osteoarthritis Cartilage. 2011 Jul 1;19(7):779–91.

12. Pareek A, Carey JL, Reardon PJ, Peterson L, Stuart MJ, Krych AJ. Long-Term Outcomes after Autologous Chondrocyte Implantation. Cartilage. 2016 Oct;7(4):298–308.

13. Ma B, Leijten JCH, Wu L, Kip M, van Blitterswijk CA, Post JN, et al. Gene expression profiling of dedifferentiated human articular chondrocytes in monolayer culture. Osteoarthritis Cartilage. 2013 Apr;21(4):599–603.

14. Wang L, Verbruggen G, Almqvist KF, Elewaut D, Broddelez C, Veys EM. Flow cytometric analysis of the human articular chondrocyte phenotype in vitro. Osteoarthritis Cartilage. 2001 Jan;9(1):73–84.

15. Diaz-Romero J, Gaillard JP, Grogan SP, Nesic D, Trub T, Mainil-Varlet P. Immunophenotypic analysis of human articular chondrocytes: changes in surface markers associated with cell expansion in monolayer culture. J Cell Physiol. 2005 Mar;202(3):731–42.

16. Hamada T, Sakai T, Hiraiwa H, Nakashima M, Ono Y, Mitsuyama H, et al. Surface markers and gene expression to characterize the differentiation of monolayer expanded human articular chondrocytes. Nagoya J Med Sci. 2013 Feb;75(1-2): 101–11.

17. Williams R, Khan IM, Richardson K, Nelson L, McCarthy HE, Analbelsi T, et al. Identification and clonal characterisation of a progenitor cell sub-population in normal human articular cartilage. PloS One. 2010;5(10):e13246.

18. Nelson L, McCarthy HE, Fairclough J, Williams R, Archer CW. Evidence of a Viable Pool of Stem Cells within Human Osteoarthritic Cartilage. Cartilage. 2014 Oct;5(4):203–14.

19. McCarthy HE, Bara JJ, Brakspear K, Singhrao SK, Archer CW. The comparison of equine articular cartilage progenitor cells and bone marrow-derived stromal cells as potential cell sources for cartilage repair in the horse. Vet J Lond Engl 1997. 2012 Jun;192(3):345–51.

20. Fellows CR, Williams R, Davies IR, Gohil K, Baird DM, Fairclough J, et al. Characterisation of a divergent progenitor cell sub-populations in human osteoarthritic cartilage: the role of telomere erosion and replicative senescence. Sci Rep [Internet]. 2017 Feb 2 [cited 2017 Feb 24];7. Available from: http://www.ncbi.nlm.nih.gov/pmc/articles/PMC5288717/

21. Dowthwaite GP, Bishop JC, Redman SN, Khan IM, Rooney P, Evans DJR, et al. The surface of articular cartilage contains a progenitor cell population. J Cell Sci. 2004 Feb 29;117(Pt 6):889–97.

22. Williams R, Khan IM, Richardson K, Nelson L, McCarthy HE, Analbelsi T, et al. Identification and Clonal Characterisation of a Progenitor Cell Sub-Population in Normal Human Articular Cartilage. PLOS ONE. 2010 Oct 14;5(10):e13246.

23. Marsano A, Millward-Sadler SJ, Salter DM, Adesida A, Hardingham T, Tognana E, et al. Differential cartilaginous tissue formation by human synovial membrane, fat pad, meniscus cells and articular chondrocytes. Osteoarthritis Cartilage. 2007 Jan 1;15(1):48–58.

24. Yang ZX, Han Z-B, Ji YR, Wang YW, Liang L, Chi Y, et al. CD106 identifies a subpopulation of mesenchymal stem cells with unique immunomodulatory properties. PloS One. 2013;8(3):e59354.

25. Kienzle G, von Kempis J. Vascular cell adhesion molecule 1 (CD106) on primary human articular chondrocytes: functional regulation of expression by cytokines and comparison with intercellular adhesion molecule 1 (CD54) and very late activation antigen 2. Arthritis Rheum. 1998 Jul;41(7):1296–305.

26. Dominici M, Le Blanc K, Mueller I, Slaper-Cortenbach I, Marini F, Krause D, et al. Minimal criteria for defining multipotent mesenchymal stromal cells. The International Society for Cellular Therapy position statement. Cytotherapy. 2006;8(4):315–7.

27. Knudson CB, Knudson W. Hyaluronan and CD44: modulators of chondrocyte metabolism. Clin Orthop. 2004 Oct;(427 Suppl):S152–162.

28. Jayasuriya CT, Chen Q. Potential benefits and limitations of utilizing chondroprogenitors in cell-based cartilage therapy. Connect Tissue Res. 2015;56(4):265–71.

29. Koelling S, Kruegel J, Irmer M, Path JR, Sadowski B, Miro X, et al. Migratory chondrogenic progenitor cells from repair tissue during the later stages of human osteoarthritis. Cell Stem Cell. 2009 Apr 3;4(4):324–35.

30. Fujita Y, Shiomi T, Yanagimoto S, Matsumoto H, Toyama Y, Okada Y. Tetraspanin CD151 is expressed in osteoarthritic cartilage and is involved in pericellular activation of pro-matrix metalloproteinase 7 in osteoarthritic chondrocytes. Arthritis Rheum. 2006 Oct;54(10):3233–43.

31. Swart GWM. Activated leukocyte cell adhesion molecule (CD166/ALCAM): developmental and mechanistic aspects of cell clustering and cell migration. Eur J Cell Biol. 2002 Jun;81(6):313–21.

32. Su X, Zuo W, Wu Z, Chen J, Wu N, Ma P, et al. CD146 as a new marker for an increased chondroprogenitor cell sub-population in the later stages of osteoarthritis. J Orthop Res Off Publ Orthop Res Soc. 2015 Jan;33(1):84–91.

33. Vinod E, Boopalan PRJVC, Sathishkumar S. Reserve or Resident Progenitors in Cartilage? Comparative Analysis of Chondrocytes versus Chondroprogenitors and Their Role in Cartilage Repair. Cartilage. 2017 Oct 1;1947603517736108.

34. Mobasheri A, Kalamegam G, Musumeci G, Batt ME. Chondrocyte and mesenchymal stem cell-based therapies for cartilage repair in osteoarthritis and related orthopaedic conditions. Maturitas. 2014 Jul;78(3):188–98.

35. Sophia Fox AJ, Bedi A, Rodeo SA. The Basic Science of Articular Cartilage. Sports Health. 2009 Nov;1(6):461–8.

36. Bernstein P, Sperling I, Corbeil D, Hempel U, Fickert S. Progenitor cells from cartilage-no osteoarthritis-grade-specific differences in stem cell marker expression. Biotechnol Prog. 2013 Feb;29(1):206–12.

37. Nakauchi H. Hematopoietic stem cells: Are they CD34-positive or CD34-negative? Nat Med. 1998 Sep;4(9):1009–10.

38. Roughley PJ. The structure and function of cartilage proteoglycans. Eur Cell Mater. 2006;12:92–101.

39. Benz K, Stippich C, Freudigmann C, Mollenhauer JA, Aicher WK. Maintenance of ‘stem cell’ features of cartilage cell sub-populations during in vitro propagation. J Transl Med. 2013;11:27.

40. Pretzel D, Linss S, Rochler S, Endres M, Kaps C, Alsalameh S, et al. Relative percentage and zonal distribution of mesenchymal progenitor cells in human osteoarthritic and normal cartilage. Arthritis Res Ther. 2011;13(2):R64.

41. Su X, Zuo W, Wu Z, Chen J, Wu N, Ma P, et al. CD146 as a new marker for an increased chondroprogenitor cell sub-population in the later stages of osteoarthritis. J Orthop Res Off Publ Orthop Res Soc. 2015 Jan;33(1):84–91.

42. Schwab KE, Hutchinson P, Gargett CE. Identification of surface markers for prospective isolation of human endometrial stromal colony-forming cells. Hum Reprod Oxf Engl. 2008 Apr;23(4): 934–43.

43. Jiang Y, Cai Y, Zhang W, Yin Z, Hu C, Tong T, et al. Human Cartilage-Derived Progenitor Cells From Committed Chondrocytes for Efficient Cartilage Repair and Regeneration. Stem Cells Transl Med. 2016 Jun;5(6):733–44.

44. McCarthy HE, Bara JJ, Brakspear K, Singhrao SK, Archer CW. The comparison of equine articular cartilage progenitor cells and bone marrow-derived stromal cells as potential cell sources for cartilage repair in the horse. Vet J Lond Engl 1997. 2012 Jun;192(3):345–51.

